# Dependence of Heart Rate Variability on Immersion in a Virtual Game and Change of Visual Content in the Flow of Virtual Reality

**DOI:** 10.1101/2021.10.06.463287

**Authors:** Vasily Pyatin, Arseny Videnin, Olga Maslova, Sergei Chaplygin, Sergey Rovnov

## Abstract

A modern person constantly changes the environment of his mental activity, moving real into an immersive environment, for example, from the surrounding reality to the information environment of a smartphone and back. This kind of transition is needed to satisfy many of the cognitive and emotional needs of people. The transition from a real physical environment to virtual reality (VR) with the help of a special headset, for example, the Oculus Rift or HTC vive, occurs less often, causing less frequent emotional state. If at the same time the emotional state of a person is investigated, then, as a rule, the manifestation of heart rate variability (HRV) is used as an indicator. However, there are relatively few studies in the literature on physiological responses using HRV during the perception of VR content. The results of such studies are extrapolated to data evoked in HRV manifestations by stimuli of real and virtual environments. We studied HRV in 55 participants while they were in the flow of VR content of different dynamics. The results were analyzed by statistically testing the hypothesis of the effect of VR immersion and the effect of transitions between realities on HRV manifestations, as well as the effect of VR flow dynamics on HRV. The results showed that the perception of the VR flow and the content transitions made in it determine the change in HRV in the form of such parameters as LF and the Baevsky Index, which can be considered as markers of immersion in VR. An increase in emotional arousal with sequential participation in virtual games in one virtual stream determines the manifestations of HRV - HR, Moda and PAR.

The results contribute to understanding the possibilities of using VR technology to recognize emotions during the transition between the natural environment and VR, as well as to determine the level of emotional arousal when immersed in the changing flow of virtual content. These studies are important for the study of the psychology of emotions in the VR flow paradigm.

## Introduction

A huge number of people in the world make a mind transition from the real-world to a smartphone and back to solve their cognitive and communication problems. [1–2]. The situation of emotional and informational transitions also occurs when using a VR helmet, which is widely used in the games industry [3], in medicine [4–5], in teaching [6–7] in virtual tourism [8–9], practicing practical skills, for example, in sociology [10], in VR cinema [11]. A relatively small number of studies to assess human physiological responses during the perception of VR content [12–14] are based on HRV. As a rule, previous studies extrapolate physiological data from being in VR with those caused by being in a real information environment. We did not find any research results among published scientific papers that investigated emotional responses to changing content directly within the visually perceived virtual stream. The meaning of our work is to study the manifestations of HRV as a reflection of the sympatho-vagus balance of the emotional process caused by the flow of information content within VR. Our study compares the manifestations of HRV during active actions in VR, as well as during passive stay in VR and outside VR; searches for markers that represent immersion in VR. The publications [12–14] show the possibility of assessing the degree of immersion in virtual game content by the manifestations of HRV. The use of this approach is due to the fact that indicators of HRV or electrical activity of the skin may be more suitable for assessing the emotional state of gamers than psychological tests. Obviously, using questionnaires, it is impossible to assess the psychophysiological state of a gamer in real time [14]. However, the issue requires further study, since the relationship between the degree of content interactivity and changes in HRV indicators is not clear, and it is also necessary to clarify the relationship of the emotional state with specific HRV parameters. In addition, no attempt has been made to separate the impact of vigorous physical activity by the player from the specific impact of immersion in VR based on HRV manifestations. In our work, we compare the manifestations of HRV during immersion in the flow of changing virtual activity.

## Methods

### Participants

The study involved fifty-five healthy young people aged 21 ± 3 years from among students of the Samara State Medical University: twenty-eight boys and twentyseven girls. Participants learned about the study from social media advertisements and from university faculty and lecturers. Participation was not intended to receive direct benefits and monetary rewards. The participants met the following inclusion criteria: did not suffer from diseases of the cardiovascular system and nervous system, did not suffer from diabetes mellitus, did not take beta-blockers or selective serotonin reuptake inhibitors. The exclusion criteria were: high anxiety, the onset of nausea when interacting with VR, a dark opaque manicure that made photoplethysmography difficult with a finger sensor.

### Ethics

The study was approved by the Bioethics Committee of the Samara State Medical University. All participants gave written informed consent to participate in the study and written consent to the processing of personal data.

### Study protocol

The VR complex HTC Vive pro and the virtual game Bow from the collection The Lab (Steam) were used. During the virtual game, the participants performed active actions, shooting from a bow at moving virtual warriors who were actively moving around the playing field with different dynamics of the process. The game frame continued either until the participant won, or until the participant lost in the game “Game over” (Fig 1). During the transition of a participant in the VR stream from active actions in the game to the task “passive observation”, the participants still remained wearing VR glasses, but they were shown a “freeze frame” in the form of a static playing space without active actions.

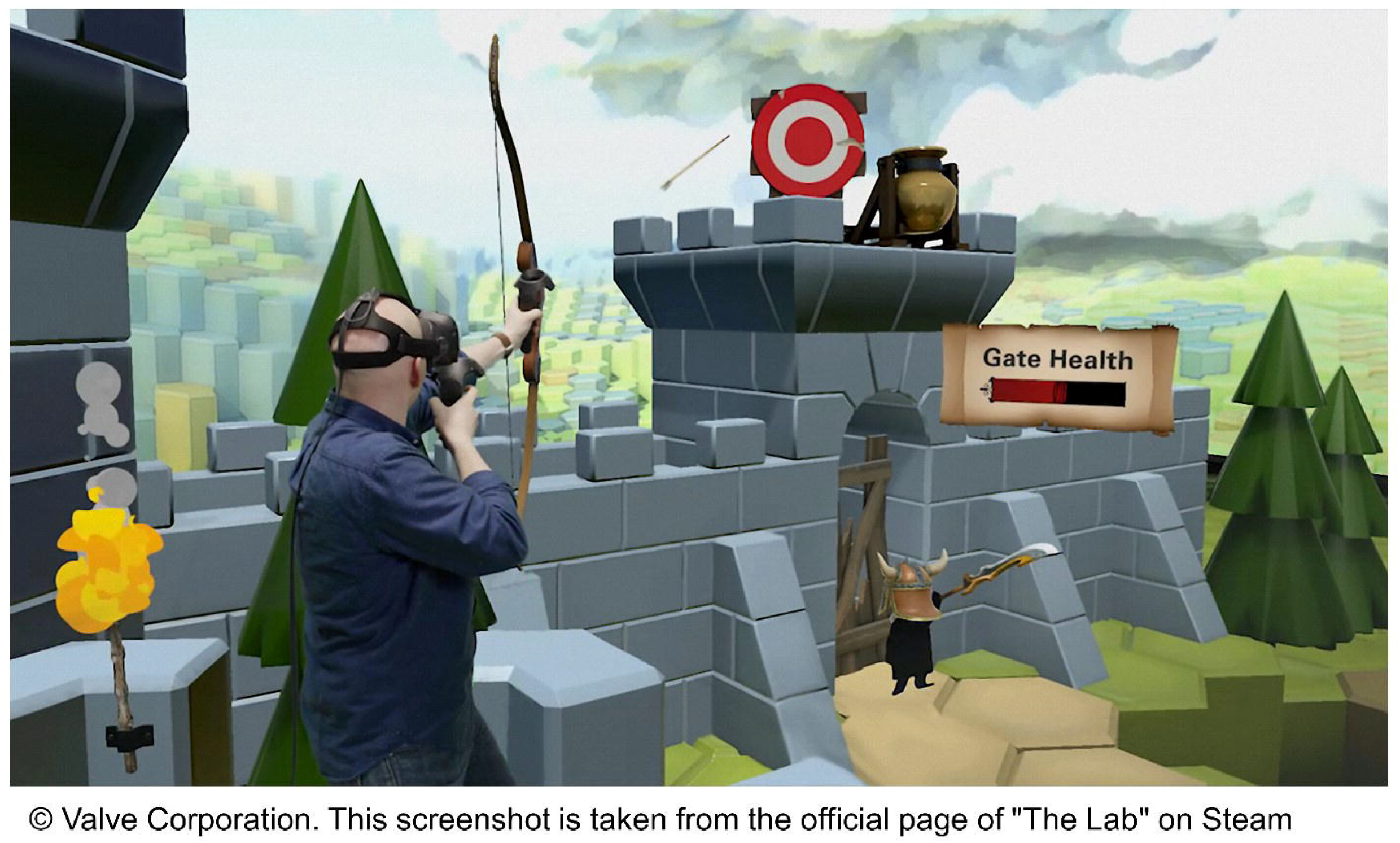

Participants were required not to smoke or drink caffeinated beverages within 3 hours of the experiment. 55 participants were randomly divided into three groups (Fig 2).

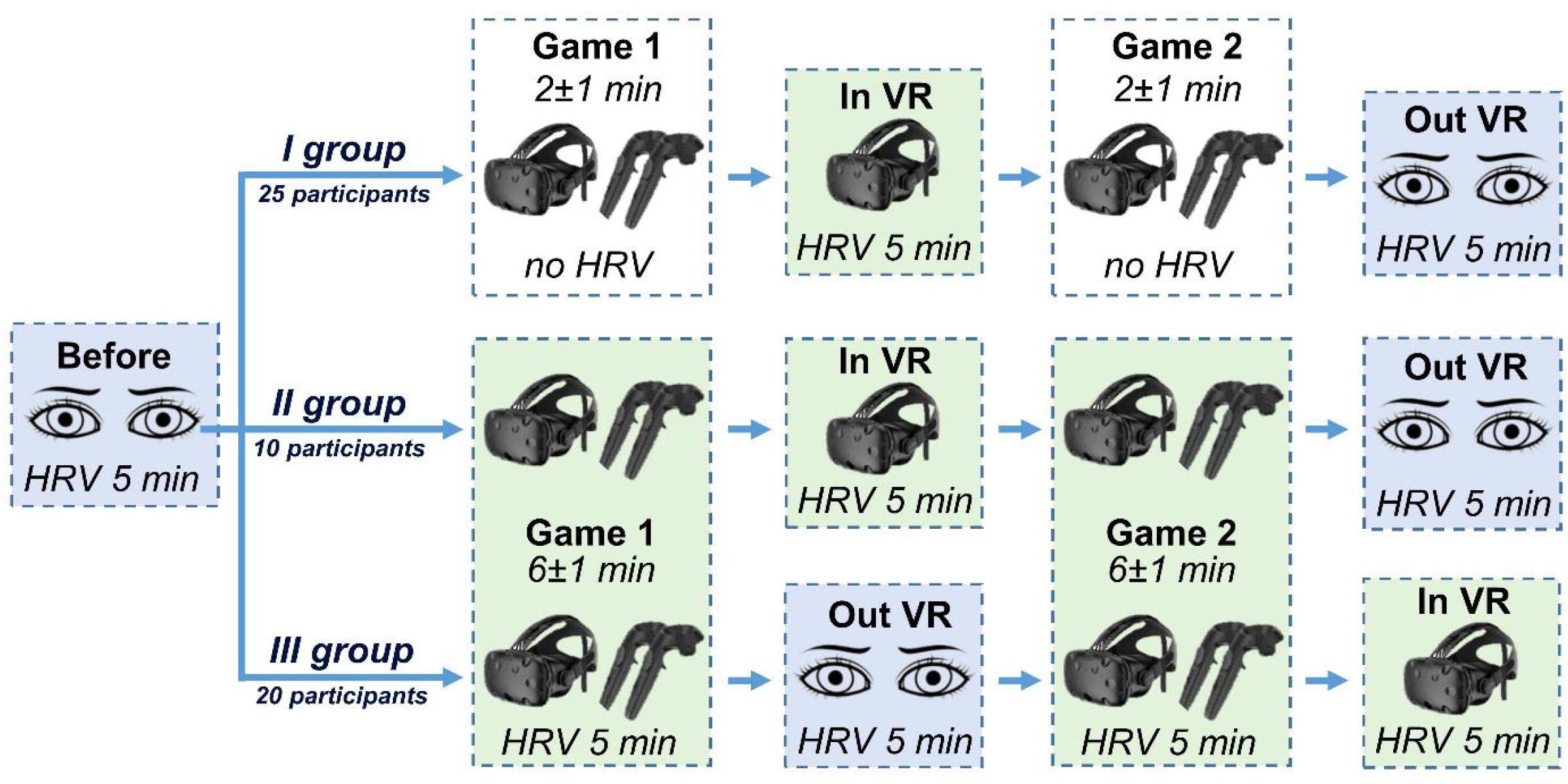

Twenty-five participants were randomly assigned to Group I. During the experiment, they recorded manifestations of HRV at the following points: at baseline (5 min, HRV “Before”); during a break between virtual games in VR (5 min, HRV “in VR”) and immediately after the completion of the second virtual game outside VR (5 min, HRV “Out VR”). The duration of each of the two virtual games was 2 ± 1 minutes and differed depending on how effectively the subjects played: the higher the efficiency, the longer the game lasted (survival game). In group I, HRV manifestations during two virtual games were not recorded due to the short time of each virtual game (2 + 1 min) and due to the fact, that, according to the European standard, HRV registration should last at least 5 minutes [15].

Ten study participants were randomly assigned to group II. In this series of experiments, HRV manifestations were recorded at all stages of the session: in the initial state (5 min, HRV “Before”), during participation in each of two virtual games (6 + 1 min, HRV “Game 1” and “Game 2”) and during passive observation of a static panorama between virtual games (5 min, HRV “in VR”) and outside VR after the completion of the second virtual game (5 min, HRV “Out VR”). The duration of the virtual game session for the participants of group II was 5-7 minutes in order to be able to register the manifestations of HRV within 5 minutes. To do this, when the game ended early, earlier than 5 minutes later, the game was restarted and the participants continued to play until they won or lost. In this case, HRV was recorded within 5 minutes from the start of the virtual game.

Twenty participants were randomly assigned to group III. As shown in Fig 2, the study protocol of this group differed from that of the study group II in that after the first virtual game, the participants took off their VR glasses and went for 5 minutes (HRV “Out VR”) into the real environment. After the second virtual game, the participants passed to passively observing the static content of the virtual scene without removing their VR glasses (5 min, HRV “in VR”). These differences in experimental protocols for group II and group III participants were undertaken to assess the effects of fatigue and virtual habituation factors on HRV manifestations. HRV was recorded using an ELOKS-01 pulse oximeter (CJSC IMC “New devices”, Russian Federation) by transmission photoplethysmography (pulsometry). When registering by this method, the structural unit of the heart rate is the time interval between the highest points of the pulse waves - the NN (NN) interval, an analogue of the RR interval of the electrocardiogram. Primary data processing in the ELOGRAPH 3.0 program made it possible to determine the values of 18 heart rate parameters using standard methods [15–16]: heart rate; NN fashion; the amplitude of the NN mode; standard deviation NN; the square root of the mean squares of the difference of consecutive NN; percentage of consecutive NN pairs differing by more than 50 ms; variation range NN; HRV index; triangular interpolation index of the histogram NN; parameters of the spectrum of NN oscillations (by the method of

Fourier transforms) in the range of very low (VLF), low (LF), and high (HF) frequencies, spectrum power (Total), parameters of the ratio LF and HF (pLF, pHF, LF / HF); the index of activity of the sympathetic (SYM) and parasympathetic (PAR) divisions of the autonomic nervous system, the index of tension of regulatory systems according to Baevsky (Baevsky index).

### Statistical analysis

Statistical processing of data, construction of tables and diagrams were carried out in Statistics 12 and Excel 2016. Due to the deviation of the distribution of the majority (70%) of the samples from the normal distribution (p0 according to the Shapiro-Wilk, Kolmogorov-Smirnov, Lilliefors criteria <0.05), the samples were described median and quartiles, the significance of differences between registrations of the same participants was calculated using the Wilcoxon test. Due to the high variability of the results due to the specificity of the HRV method and did not allow to achieve homogeneity of the groups for a given sample size, the study of intergroup differences was not carried out. The paper presents the values of only those HRV parameters that significantly differed when comparing comparable registrations: between passive observation registrations after a virtual game and between registrations during a virtual game.

## Results

Table 1 shows the manifestations of HRV of participants in all three groups after the termination of the virtual game: 1) the transition from the virtual game to passive observation in VR (Group I-III); 2) the transition from a virtual game to the surrounding natural reality (Group I-III).

**Tabl 1.** Significant differences between the states of passive observation

|  |  | Bayevsky<br>index, c.u. | Low frequency,<br>ms<sup>2</sup> | Total power, ms<sup>2</sup> |
| --- | --- | --- | --- | --- |
| In VR –<br>out VR | M (Q<sub>1</sub>; Q<sub>3</sub>) | 6 (-12; 33) | -426 (-1765; 398) | -372 (-3799; 1406) |
|  | p0 (Wilcoxon test) | 0,031 | 0,025 | 0,145 |
| In VR –<br>before | M (Q<sub>1</sub>; Q<sub>3</sub>) | 7 (-13; 35) | -407 (-1347; 676) | -1203 (-3611; 1640) |
|  | p0 (Wilcoxon test) | 0,081 | 0,092 | 0,033 |
| Out VR<br>–<br>before | M (Q<sub>1</sub>; Q<sub>3</sub>) | 1 (-27; 33) | 297 (-927; 1319) | -353 (-3087; 2117) |
|  | p0 (Wilcoxon test) | 0,645 | 0,530 | 0,487 |

In this work, we compared the manifestations of HRV in groups I-III between HRV data during passive observation of the VR scene after the end of the virtual game (Fig. 3, HRV “In VR”) and the data recorded after the end of the virtual game and removal of the VR helmet (Fig. 3, HRV “Out VR”). We found that when passively wearing a VR helmet after a virtual game in comparison with participants leaving VR into the environment, there is a significant decrease (Fig 3) in the power of low-frequency fluctuations in the duration of the NN-interval (LF) (Valid N = 55; T = 503; Z = 2.237; p0 = 0.025) lower by 426 (1765; −398) ms2. At the same time, the value of the stress index of regulatory systems (IB) was significantly (Valid N = 54; T = 492; Z = 2.157; p0 = 0.031) higher by 6 (−12; 33) conventional units. These differences in the manifestations of HRV, identified according to the paradigm of the protocol of our study, are markers of the participants’ degree of immersion in VR during the game. It should be emphasized that the state “in VR” and “Out VR”, except for the factor of immersion in VR, did not differ in any way, since the design of the protocol allows excluding the influence of factors of addiction and fatigue as a result of playing in VR, since in the study protocol (group III, N = 20) the experimental conditions were changed in their sequence. So, in the protocol of experiments for groups I and II (N = 35), after the first virtual game, the participants went into the “in VR” state, and after the second virtual game, into the “Out VR” state. On the contrary, in the experiment protocol for group III (N = 20), the first virtual game was followed by the “Out VR” state, and the second virtual game was followed by the “in VR” state. Therefore, it can be assumed that a decrease in LF values and an increase in IB are correlates of the psychoemotional state of immersion in VR. These changes in the LF and IB parameters indicate the centralization of the regulation of sympatho-vagus processes in VR.

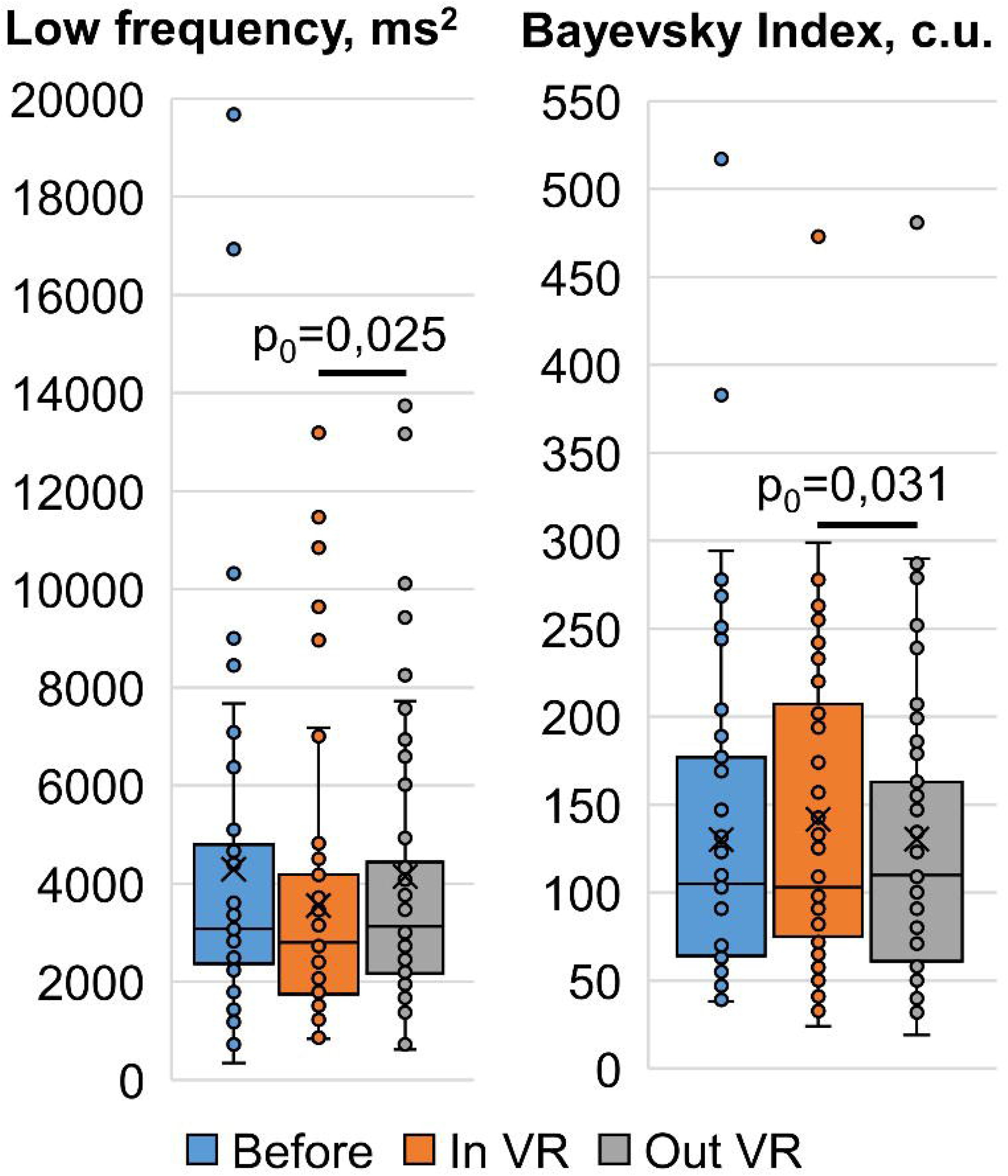

Analysis of HRV manifestations within the VR content stream (Game 1 and Game 2) revealed such correlates of the participants’ emotional state as HR, Moda, and PAR responses. For these manifestations of HRV, we compared them during the second virtual game with those during the first virtual game (Fig 4). So, during the second virtual game relative to the first in the general flow of VR, the manifestations of HRV were as follows: HR was significantly (Valid N = 28; T = 94; Z = 2.482; p0 = 0.013) higher by 2 (−1; 6) b.p.m.; Moda - significantly (Valid N = 26; T = 96; Z = 2.019; p0 = 0.043) lower by 15 (30; −8) ms, and PAR index - significantly (Valid N = 28; T = 105.5; Z = 2.220; p0 = 0.026) 1 (4; −1) conventional units lower. Note that the content complexity of the first and second virtual games was identical. It can be assumed on the basis of these manifestations of HRV that changes in HR, Moda and PAR reflect a higher level of emotional arousal of the participants during the second virtual game, and these HRV parameters can be attributed to the objective metrics of studying the vegetative components of emotions.

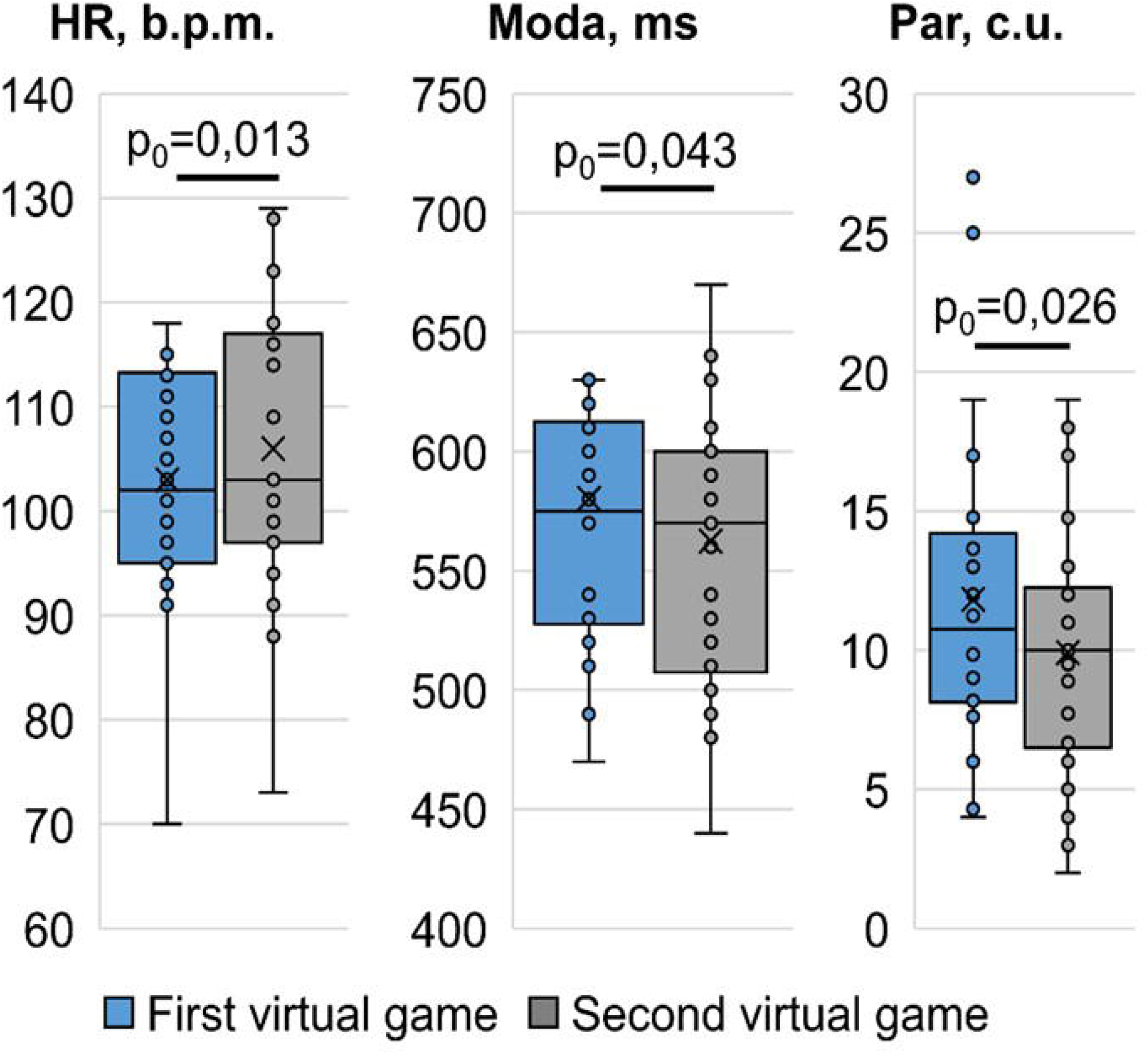

## Discussion

A less common informational transition is dynamic transactions between real-world and VR using a dedicated headset like the Oculus Rift or HTC vive. In our work, we studied the manifestations of HRV, as a reflection of sympatho-vagus balance, during transitions from a real environment to VR using HTC vive glasses, and also investigated the manifestations of HRV when changing VR content, when participants are immersed in a virtual game, performing active game actions in it. This made it possible to objectively assess the degree of immersion of participants in VR according to HRV data. A similar paradigm can be applied to the study of emotions with a sequential stream of virtual scenes of different content and the number of presented virtual frames. Currently, according to [17], the evolution of scientific interest in VR and emotions caused by immersion in VR is growing exponentially at the rate of more than 200 publications per year. This is due to the wider use of VR in recent years in areas such as rehabilitation, neurosurgery and therapy [18].

The research carried out under our hypothesis is based on the assumption that VR is likely to change areas of human activity such as psychological research, watching films, for example, and the way films are produced, promoted and sold [11]. Currently, the innovative potential of VR content production is increasing, the production ecology of its scripts is cultivated and high quality VR content is provided. Emotions play a key role in our daily life, so understanding and recognizing emotional responses is critical to research in an emerging technology like VR. Immersive VR, which allows researchers to simulate the environment in a controlled laboratory setting with a high level of presence and interactivity, is becoming increasingly popular in emotion research [20]. In recent years, VR has also become more popular in the scientific and commercial contexts [17], and the general level of VR applications includes a wide range of activities - games, learning, education, healthcare, marketing and more. Studies have used a variety of signals such as voice, face, neuroimaging and physiological signals to recognize a person’s emotional responses when immersed in VR [19–20]. In particular, HRV manifestations are effective markers in recognizing emotional responses to virtual content.

However, the number of known published studies in this area is limited [12–14]. At the same time, they are all aimed at extrapolating the differences in the HRV reaction between the perception of the content of the real environment and VR. In our work, for the first time, we studied the features of the psychoemotional state according to HRV data during the transition of study participants from the real environment to VR and vice versa. Within the framework of such an experiment design, HRV indicators of subjects’ immersion in a virtual game were identified, but also an increase in emotional stress with active participation in a virtual stream over a longer period of time in a virtual game. Further studies of the dynamics of the emotional state are required within the framework of the outlined paradigm of the VR content flow. This can contribute to the development of environmental modeling in a laboratory, where a high level of sense of presence and interactivity is created, as well as psychological and somatic parameters of response to this environment are controlled, which is now in great demand for the increasingly popular field of emotion research. The paradigm is of interest for the development of various psychological theories that contribute to understanding how the virtual gaming aspect can change the emotional state of the players.

## Conclusions

Expanding exponentially the cognitive and emotional transition between real and virtual environments using VR glasses. In our study, we tested the hypothesis of the possibility of assessing the degree of immersion in VR and the dynamics of VR content when immersed in a virtual game based on the manifestations of HRV. The design of the research protocol allowed us to show that the manifestations of HRV immersion in VR are LF and the Baevsky index, and the degree of emotional arousal arising in the flow of a virtual game (HR, Moda, PAR). The research results are applicable to study the psychology of emotions in VR based on the manifestations of HRV, and can also be applied in the development of the philosophy of VR cinema.

## Author Contributions

**Conceptualization**: Vasily Pyatin, Arseny Videnin, Olga Maslova, Sergei Chaplygin

**Data curation:** Vasily Pyatin, Arseny Videnin, Olga Maslova, Sergey Rovnov

**Formal analysis**: Vasily Pyatin, Arseny Videnin, Sergey Rovnov

**Investigation**: Arseny Videnin, Olga Maslova, Sergey Rovnov

**Methodology:** Vasily Pyatin, Arseny Videnin, Sergey Chapyigin

**Supervision:** Vasily Pyatin, Olga Maslova,

**Writing – original draft**: Vasily Pyatin, Arseny Videnin, Olga Maslova, Sergei Chaplygin

**Writing – review & editing**: Vasily Pyatin, Arseny Videnin, Olga Maslova, Sergey Rovnov

